# Riding the Cell Jamming Boundary: Geometry, Topology, and Phase of Human Corneal Endothelium

**DOI:** 10.1101/236406

**Authors:** Nigel H. Brookes

## Abstract

It is important to assess the viability of eye-banked corneas prior to transplantation due to inherent senescence and known loss of endothelial cells during surgical manipulation. Corneal endothelial cells have a complex basal and paracellular shape making them challenging to accurately measure, particularly in oedematous *ex vivo* tissue. This study used calibrated centroidal Voronoi Diagrams to segment cells in images of these human corneas, in order to characterize endothelial geometry, topology, and phase.

Hexagonal cells dominated the endothelia, with most comprised of five different pleomorphs exhibiting self-similar topological coarsening through most of the endothelial cell density range. There was a linear relationship between cell size and shape, though cells with greater than six sides were present in larger proportions than cells with less. Hexagonal cell regularity was stable and largely independent of density.

Cell and tissue phase was also examined, using the cell shape index relative to the recently discovered ‘cell jamming’ phase transition boundary. Images showed fluid endothelia with a range of shape indices spanning the boundary, independent of density but dependent on hexagonal regularity. The cells showed a bimodal distribution centred at the boundary, with the largest proportion of cells on the fluid side. A shoulder at the boundary suggested phase switching via shape transformation across the energy barrier, with cells either side having distinctly different size and shape characteristics. Regular hexagonal cells were closest to the boundary.

This study showed the corneal endothelium acts as a glassy viscous foam characterized by well-established physical laws. Endothelial cell death transiently and locally increases cell fluidity, which is subsequently arrested by jamming of the pleomorphically diverse cell collective, via rearrangement and shape change of a small proportion of cells, which become locked in place by their neighbours and maintain endothelial function with little energy expenditure.

## Abbreviations

CJB: Cell Jamming Boundary
CV: Coefficient of Variation
CVT: Centroidal Voronoi Tessellation
ECD: Endothelial Cell Density
SD: Standard Deviation
SEM: Standard Error of the Mean
SI: Shape Index
VD: Voronoi Diagram

## 1. Introduction

### 1.1 Microscopy and Image Analysis

Recent immunohistochemical studies have confirmed that corneal endothelial cells have a complex shape, comprised of foam-like apical pole coupled to a complex interdigitating stellate basal pole that is tightly anchored to Descemet’s membrane [Harrison *et al*., 2016; He *et al*., 2016]. Traditional examination of only the apical view *in vivo* with a specular or confocal microscope can only provide limited information about the endothelium, or the cornea as a whole.

The *ex vivo* corneas examined in this study were organ-culture stored and assessed using hypotonic sucrose to swell and make their intercellular borders visible [He *et al.*, 2016; Kirk and Hassard, 1969; Pels, 1997; Sperling, 1980]. While this makes the complex basal morphology more evident, it also makes morphometric analysis more difficult using common edge-based segmentation techniques, so every segmentation boundary is an approximation. This has led to the alternative approach used in this study, where point clouds of cell centroids were used to generate calibrated Voronoi Diagrams (VD), which define the region occupied by each cell and their borders [Brookes, 2017a]. Endothelial polymegathism and pleomorphism was measured using the size and shape of these regions, facilitating the large-scale examination of corneal endothelial geometry, topology and phase [Brookes, 2017b].

### 1.2 Cell Density and Shape

The viability of an eye-banked cornea for transplantation is usually quantified by microscopically determining endothelial cell density (ECD) [Wilson and Bourne, 1989], which declines constantly throughout life [Wilson and Roper-Hall, 1982]. While the density of the endothelial molecular pumps has been shown to remain remarkably constant in order to maintain normal corneal thickness and transparency [Geroski *et al*., 1985], corneal decompensation and failure can result when the ECD drops to very low levels [Mishima, 1982, Nishimura, 1999]. This is a problem as much corneal tissue stored for transplant is from older donors [Cunningham *et al*., 2012], and transplantation surgery inevitably causes cell loss, particularly popular lamellar procedures such as Descemet’s Stripping Endothelial Keratoplasty [Price and Price, 2008].

This steady decrease in ECD throughout life is due to the inherent senescence of the endothelial cells, which *in vivo* are arrested in the G1-phase of the cell cycle, and dead cells are not replaced. Three mechanisms have previously been identified that contribute to this senescence: cell–cell contact-dependent inhibition correlated to the time endothelial cell division ceases and mature cell contacts form, lack of effective growth factor stimulation, and TGF-β2 suppression of the S-phase of the cell cycle [Joyce, 2005; Joyce, 2012].

Beyond ECD, the polymegathism and pleomorphism of the endothelial cells was recognized in early studies to also affect corneal viability. For example, corneal endothelia from diabetic patients showed abnormalities in cell size, shape, and arrangement, suggesting these may be early indicators of endothelial distress [Rao, 1985; Shultz *et al*., 1984], and the decline in ECD and the proportion of hexagonal cells after corneal transplantation appeared to affect the development of late endothelial failure [Bourne, 2001].

During early gestation, there is a rapid increase in the density of endothelial cells to about 16,000 cells/mm^2^, but as gestation proceeds the corneal diameter increases and this density steadily decreases. At birth, the corneal endothelium was long assumed to be a uniform monolayer comprised mostly of hexagonal cells, but more detailed analysis has shown that while it has a cell density of 4000-6000 cells/mm^2^[Elbaz *et al*., 2017; Ko *et al*., 2001] there is also a high level of pleomorphism [Doughty, 1989; Müller *et al*., 2000]. Cell density values in children are significantly correlated to corneal diameter [Elbaz *et al*., 2017; Müller and Doughty, 2002; Murphy *et al*., 1984]. This trend continues through life, with a steady decrease in hexagonal cells, and a corresponding increase in pentagonal and heptagonal cells [Yee *et al*., 1985]. A study of corneal endothelia over 100 years of age found a mean cell density of 923 cells/mm^2^ comprised of about 55% hexagonal cells [Ruusuvaara *et al*., 2015]. There are also significant differences in cell density and percentage of hexagonal cells in the central, paracentral, and peripheral endothelium [Amann *et al*., 2003; Schimmelpfennig, 1984].

As a tessellated matrix of perfectly regular hexagons can only tile a flat plane, the curved posterior corneal surface must have a more complex pleomorphic composition from the outset. Tessellation of negative curvatures tends to increase the proportion of cells with more than six sides, compared to cells with less [Weaire and Rivier, 1984].

A variety of statistics have been used to characterize the composition and disorder of the corneal endothelium. The most common are the percentage of hexagonal cells and the Coefficient of Variation (CV) of the cell size, however both these are problematic. Large CV values may be due to disproportionately smaller or larger cells of unknown polygonality [Doughty, 1990; Doughty, 2013], and the percentage of hexagonal cells provides no information on their symmetry or regularity [Doughty, 1992]. The calculation of CV of the cell area is programmed into the analysis software in many commercial specular microscopes and continues to appear in the cornea literature.

Alternative measures also appearing in the literature include the figure coefficient (or shape factor), shape deviation index, skewness index, hexagon deviation index [Alanko, 1983; Doughty, 1989; Doughty, 1992], area-side plots [Dilts and Doughty, 1996; Doughty and Dilts, 1994], and angular regularity [da Fontoura Costa, 2006], but none of these have been widely used. The area-side plots are useful as they show a linear relationship between the cell area and the number of cell sides, as long as the sample size is large enough [Doughty, 1998a]. This is a demonstration of the Lewis law, where the average area of an *n*-sided cell in a two-dimensional lattice increases linearly with *n*, so small cells tends to have fewer sides [Lewis, 1926]. This law has general application in many other fields beyond corneal endothelial architecture [Chiu, 1995].

In addition to the Lewis law a number of other factors also determine the relationship between cell polygonality and size/density in a two-dimensional tessellated lattice. In general it is also defined by Euler’s formula in two dimensions (*n*_*cells*_ – *n*_*edges*_ + *n*_*vertices*_ = *Euler constant*; in a tissue with a large number of cells the average number of neighbours of a cell will be six) [Cantat *et al*., 2013], the Aboav–Weaire law (cells with a higher number of sides tend to have few-sided neighbours and *vice versa*), and the von Neumann-Mullins relation (a cell with more than six sides grows and a cell with less than six sides shrinks, with equilibrium being obtained by hexagonal cells). Much of this understanding has been developed through the study of analogous non-biological systems, such as foams and mathematical modelling [Cantat *et al*., 2013; Chiu, 1995; MacPherson and Srolovitz, 2007; Sánchez-Gutiérrez *etal*, 2016; Weaire and Rivier, 1984; Wörner *et al*., 2011].

### 1.3 Tissue Organization

Tissue homeostasis occurs through the interplay between a robust and stable architecture that can resist stress, with one exhibiting sufficient plasticity to allow remodelling to maintain tissue function [Lecuit and Lenne, 2007]. The basic laws of organization of groups of cells in a tissue were first identified by Plateau in 1873, and the apical surface of the endothelial cells will tend toward equilibrium of faces (smooth surfaces with a constant mean curvature determined by the Young-Laplace law) and edges (surfaces always meet in threes at 120° angles) [Cantat *et al*., 2013]. At equilibrium, the tissue is at an energy minimum and cells tend to aggregate in clusters, where the surface area in contact with the surrounding environment is minimized [Lecuit and Lenne, 2007]. In the *minimal surface model* this is the point where the total length of all cell edges is at a minimum and is determined solely by the cell geometry [Weaire and Rivier, 1984], though Plateau’s laws have been shown to break down in foams as the liquid fraction increases [Drencken and Hutzler, 2015]. *in vivo*, the cell geometry is further determined by both intracellular and extracellular mechanics, including cell-to-cell and cell-to-substrate adhesion, cytoskeleton, surface tension and osmotic swelling, which all act as energy barriers preventing the system from relaxing to the global minimum of energy.

Cellular dynamics have been modelled in many tissues including corneal endothelium [Honda *et al*., 1982], vascular endothelium [Vitorino *et al*., 2011], and keratinocyte monolayers [Bock, 2010]. Cells autonomously migrate within the monolayer and turn in response to mechanical cues resulting from adhesive, drag, repulsive, and directed steering interactions with neighbouring cells. Intracellularly, actin-based protrusions such as filopodia and lamellipodia have long been accepted as key constituents in these processes, together with cellular blebbing in areas of weak membrane-cytoskeleton attachment [Charras *et al*., 2008; Norman *et al*., 2010; Norman *et al*., 2011; Reinhart-King *et al*., 2005]. There is a tendency for the local migration at the *mesoscale* to follow the local orientation of maximum principal stress, called *plithotaxis* [Pegorano et al., 2016]. This mechanism of collective cell guidance is an emergent property that requires cooperativity of mechanical stresses across many cell-to-cell junctions [Trepat and Fredburg, 2011; Sadati 2013]. In contrast, individual cell death as in the corneal endothelium causes neighbouring cells to exert a traction stress aligned toward the void, via a phenomenon called *kenotaxis*[Kim *et al*., 2013; Park *et al*., 2016]. This is also seen on larger scales during wound healing [Honda *et al*., 1982].

Changes in a range of tissues that involve remodelling of cell–cell contacts have been shown to be accompanied by an increase in (Shannon) entropy in the form of both geometric and topological disorder, by progressively shifting a mosaic from a hexagonal arrangement (the VD generated from a perfectly triangular lattice of centres) to a perturbed one with increased polymegathism and pleomorphism [Lecuit and Lenne, 2007, Mason *et al*., 2012, Weaire and Rivier, 1984]. While this final organization may be irregular it is not random [Doughty, 1998b].

As the cornea ages, every endothelial cell death transiently increases tissue entropy and fluidity, forcing cells to rearrange in order to maintain endothelial function. In the *vertex model*, a system that has been used extensively to model monolayers, these cell neighbour exchanges are known as T1 processes or transitions that occur via successive boundary shortening procedures. Cell death occurs as a series of T1 processes followed by a T2 process removing the last part of the cell [Honda and Nagai, 2015, Cantat et al., 2013; Weaire and Rivier, 1984]. These initially *kenotactic* rearrangements occur in response to a non-equilibrium state, causing tissue relaxation via von Neumann-Mullins coarsening, at a rate controlled by the amount of energy required to execute them, and the degree of collective cell fluidity [Bi *et al*., 2014]. These processes maintain Plateau’s laws and Euler’s formula, and also leave the topological charge of the tissue averaging zero [Cantat *et al*., 2013].

Much of the physical tissue properties are determined by the geometry and topology of the cells comprising it. A cell’s ability to move within the tissue is topologically constrained by its neighbours, and the number and arrangement of these neighbours in turn define its geometry. Recently a ‘cell jamming’ phase transition boundary (solid-fluid; CJB) that characterizes this glassy collective cell dynamic has been discovered in the *vertex model*, and confirmed in a Voronoi model [Bi *et al*., 2016]. It can be measured with the cell shape index (SI), a dimensionless parameter that is the ratio of the cell perimeter to the square root of the cell area, that effectively encodes the physical properties of each cell such as cell–cell adhesion and cortical tension [Bi *et al*., 2015]. SI’s higher than 3.81 define a fluid cell collective, and below, a solid (jammed) one. For example, in cancer biology this parameter can differentiate between a benign (solid) and a metastasizing (fluid) tumour, essentially encoding the epithelial-mesenchymal transition [Haeger *et al*., 2014]. Similarly, cultures of asthmatic cells from the airway epithelium retain their fluidity much longer than non-asthmatic cells [Park *et al*., 2015, Park *et al*., 2016]. The phase dynamics of the human corneal endothelium are currently unknown.

### 1.4 Summary

This study extends the dataset generated previously [Brookes, 2017a] with the additional calculation of SI for each cell, analysed in relation to cell density, size, and shape, in order to characterize endothelial tissue geometry, topology, and phase in *ex vivo* human organ-culture-stored corneas [Brookes, 2017b].

## 2. Methods

The dataset used in this study was further developed from one described previously [Brookes, 2017a]. Measurements of 354,998 cells were made from 2000 routine endothelial assessment images, taken after organ-culture storage but prior to release for transplantation, from a sequential series of 678 corneas from 351 donors, that were collected with consent by the New Zealand National Eye Bank over several years [Brookes, 2017b]. Images were analysed using custom MacOS software, with manually marked cell centroids within a fixed frame or overlapping right or bottom borders. Centroids of cells outside the frame were automatically detected, and the full point cloud used to generate a calibrated VD for the entire image field. These Voronoi ‘cells’ mapped to the observed endothelial cell borders and were used to calculate cell area and shape. Detailed statistics examined in this current study were:

*Polymegathism*

- ECD of each image (cells/mm^2^)
- Coefficient of variation (CV; standard deviation/mean) of the cell area in each image
- Cell area of all cells (μm^2^)
- Cell area of hexagonal cells (μm^2^)

*Pleomorphism*

- 4-9 sides mean percentage area in each image
- Mean number of pleomorphs (different polygonal shapes in the image)
- Mean number of cell sides in each image
- CV of the number of cell sides in each image
- Number of sides of all cells
- SI of all cells (cell perimeter/√cell area)
- Regularity of hexagonal cells (percentage difference in size ratio from a regular hexagon with short/long side diameter ratio of √3/2 = 0.866)

Tabulated size and shape data for all images and cells was transferred via tab-delimited text files into Microsoft Excel for analysis. For each image, ECD data was ranked, grouped, and the mean and standard error of the mean calculated for each corresponding group of shape data (pleomorph percentage areas, hexagonal regularity, number of pleomorphs, number of sides, coefficient of variation of the cell area and number of sides, SI). For each cell, cell area was ranked, grouped, and the mean, standard error of the mean (SEM), and standard deviation (SD) calculated for each corresponding group of shape data (number of sides, SI). For each hexagonal cell, hexagonal cell area was ranked, grouped, and the mean, SEM, and SD calculated for each corresponding group of shape data (hexagonal regularity, SI). The normality and skewness of the resulting data distributions were determined using SPSS [IBM Corp. Released 2011. IBM SPSS Statistics for Macintosh, Version 20.0. Armonk, NY: IBM Corp.], and finally all data plotted using Prism [Version 6 for Mac OS X, Graphpad Software, Inc., USA].

## 3. Results

### 3.1 Images

The image ECD data [7] was found to have a non-normal distribution (1-sample Kolomogorov-Smirnov test; p = 0) skewed negatively (−0.695), as plotted in Graph 1.

Image Mean ± SEM shape data (pleomorph 4-9 percentage areas, number of pleomorphs, number of sides, CV of the number of sides and mean cell area), was plotted against the full range of ECD values in Graphs 2- 5. Data from the single outlier image with mean density greater than 4500 cells/mm^2^ was not plotted. Graphs 3-5 were plotted using a limited y-axis subset to illustrate more subtle trends.

### 3.2 Cells

The cell area and number of sides data from the both all 354,998 cells and the subset of 182,344 hexagonal cells was ordered and grouped according to the relationship: cell area (μm^2^) = 1,000,000 ÷ ECD (cells/mm^2^) [7]. The area data of both groups of cells had a non-normal distribution (1-sample Kolomogorov-Smirnov test; p = 0) and positive skew (all cells: 2.241; hexagonal cells: 2.298). The cell area distributions were plotted in Graph 6. The cell area grouping and ordering in this graph was arranged to directly compare with the ECD grouping in Graphs 1-5.

The relationship between cell area and number of sides was plotted in Graph 7 as a line graph using a limited y-axis for comparison to Doughty’s data [Doughty 1998b], and a linear regression line calculated (y = 41.24x + 105.7; r^2^= 0.997; p < 0.0001).

The regularity of the subset of hexagonal cells relative to their cell area was plotted in Graph 8, which has a limited y-axis subset to illustrate more subtle trends. Again, the cell area grouping and ordering in this graph was arranged to directly compare with the ECD grouping in Graphs 1-5.

### 3.3 Shape Index

The image SI Mean ± SD = 3.87 ± 0.02 (Range: 3.81 – 3.93). The distribution of the mean image SI is plotted in Graph 9.

The relationship of the cell SI to mean image ECD and % area of hexagonal cells was plotted in graphs 10-11. The dotted line at a SI of 3.81 in these graphs represents the CJB [64]. Shape indices above this line indicate collective cell fluidity. The SD was plotted here to show the spread of cell SI values in relation to the CJB. Graphs 10-11 were plotted using a limited y-axis subset to illustrate more subtle trends.

The SI distribution at the cell level was plotted in Graph 12. The cell SI Mean ± SD = 3.87 ± 0.09 (Range: 3.65 – 4.72).

The cell populations under at the data points at the bimodal peaks either side of the CJB in Graph 1 were further analysed, and results summarized in Table 1 and Graphs 13-14. The cell area grouping and ordering in Graph 13 was also arranged to directly compare with the ECD groupings in other graphs.

**Table 1.**
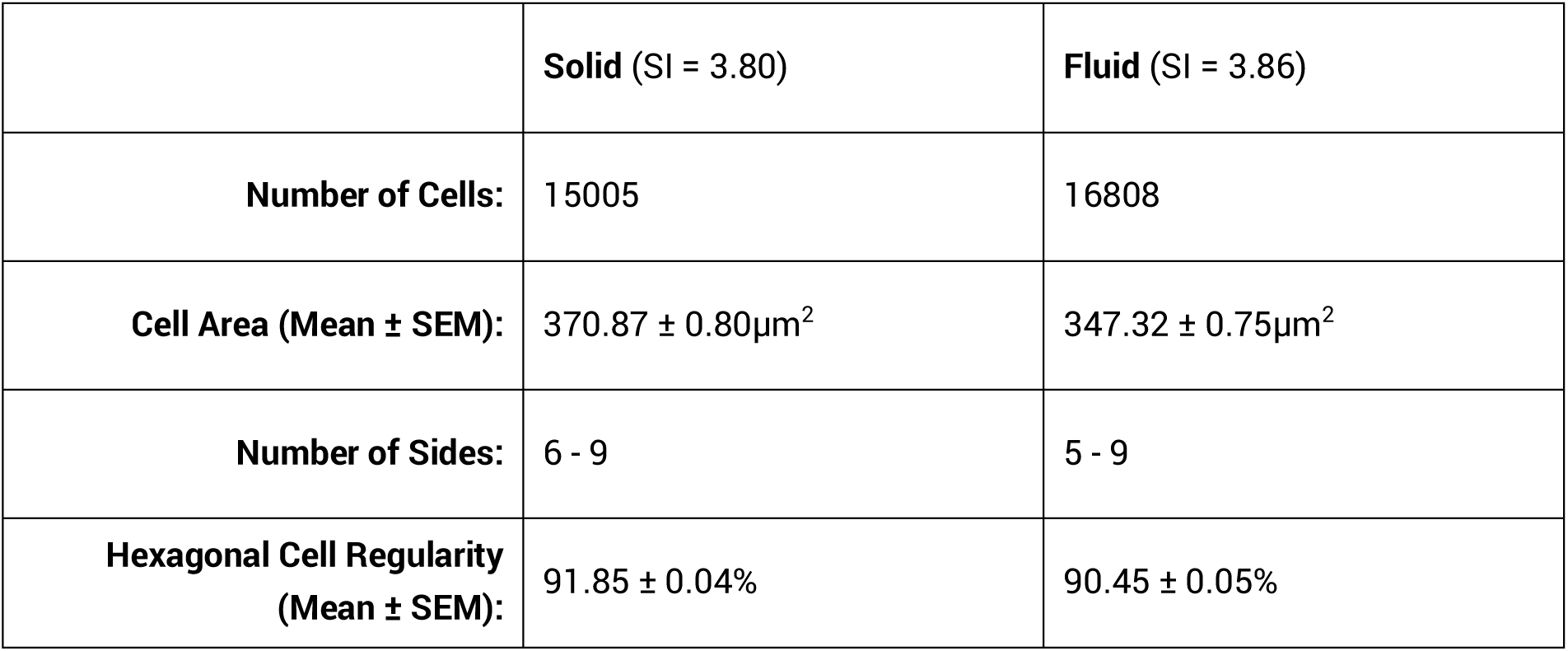
Comparative analysis of cells adjacent to the CJB.

The relationship of the SI to the hexagonal cell regularity, number of cell sides, and cell area was plotted in Graphs 15-17. The SI of the hexagonal cells in Graph 15 was sorted according to hexagonal cell regularity. The SD was again plotted to show the spread of cell SI values in relation to the CJB. These graphs were also plotted using a limited y-axis to illustrate more subtle trends.

## 4 Discussion

### 4.1 Data Collection

While the corneas used in this study were stored *ex vivo* in organ-culture, they were otherwise considered normal tissue that was almost all transplanted soon after imaging.

Swelling the intercellular spaces using hypotonic sucrose showed the convoluted basal and paracellular aspects of the cells in sharp relief using phase-contrast microscopy. The endothelial apical pole was also osmotically distorted, distinctly different from its *in vivo* appearance. These oedematous organ-cultured corneas had a folded Descemet’s membrane so the endothelium shifted in and out of the focal plane, adding further complexity that made accurately segmenting images into measurable cells very challenging. There was nevertheless an observable mapping between the endothelial cell shapes and the computed VD, suggesting analogous spatial optimization processes.

### 4.2 Endothelial Geometry and Topology

In this study, the image ECD distribution was found to be not normal but skewed slightly toward higher densities (Graph 1). Likewise, cell areas for both all cells and the subset of hexagonal cells were skewed toward smaller cell areas (Graph 6).

**Graphs 1-5.**
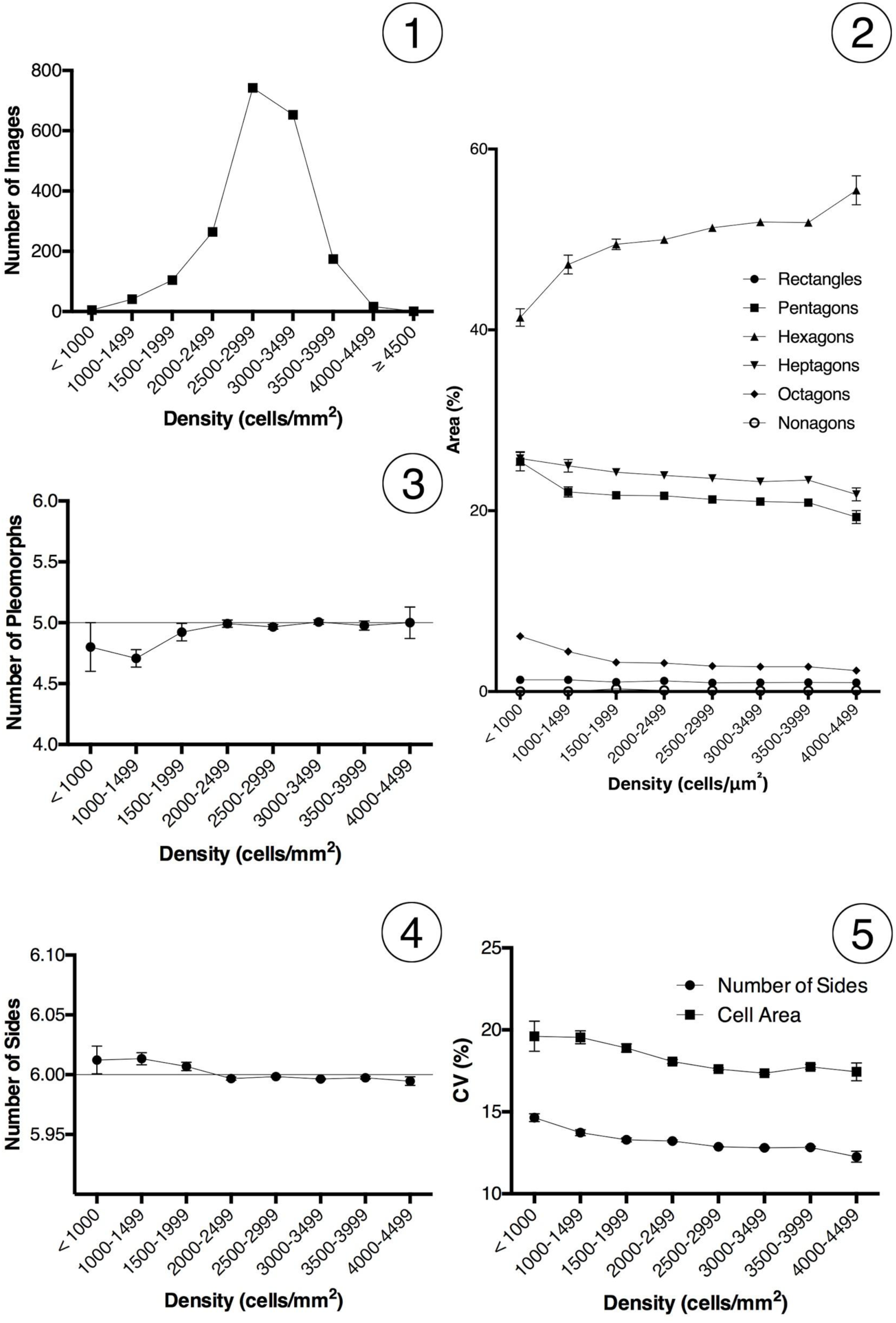
(1) Image ECD Distribution. (2) Image ECD *vs.* Image Pleomorph % Area. (3) Image ECD *vs.* Image Number of Pleomorphs. (4) Image ECD *vs.* Image Number of Sides. (5) Image ECD *vs.* Image CV of the Number of Sides / Cell Area. Graphs 2-5: Mean ± SEM.

**Graphs 6-11.**
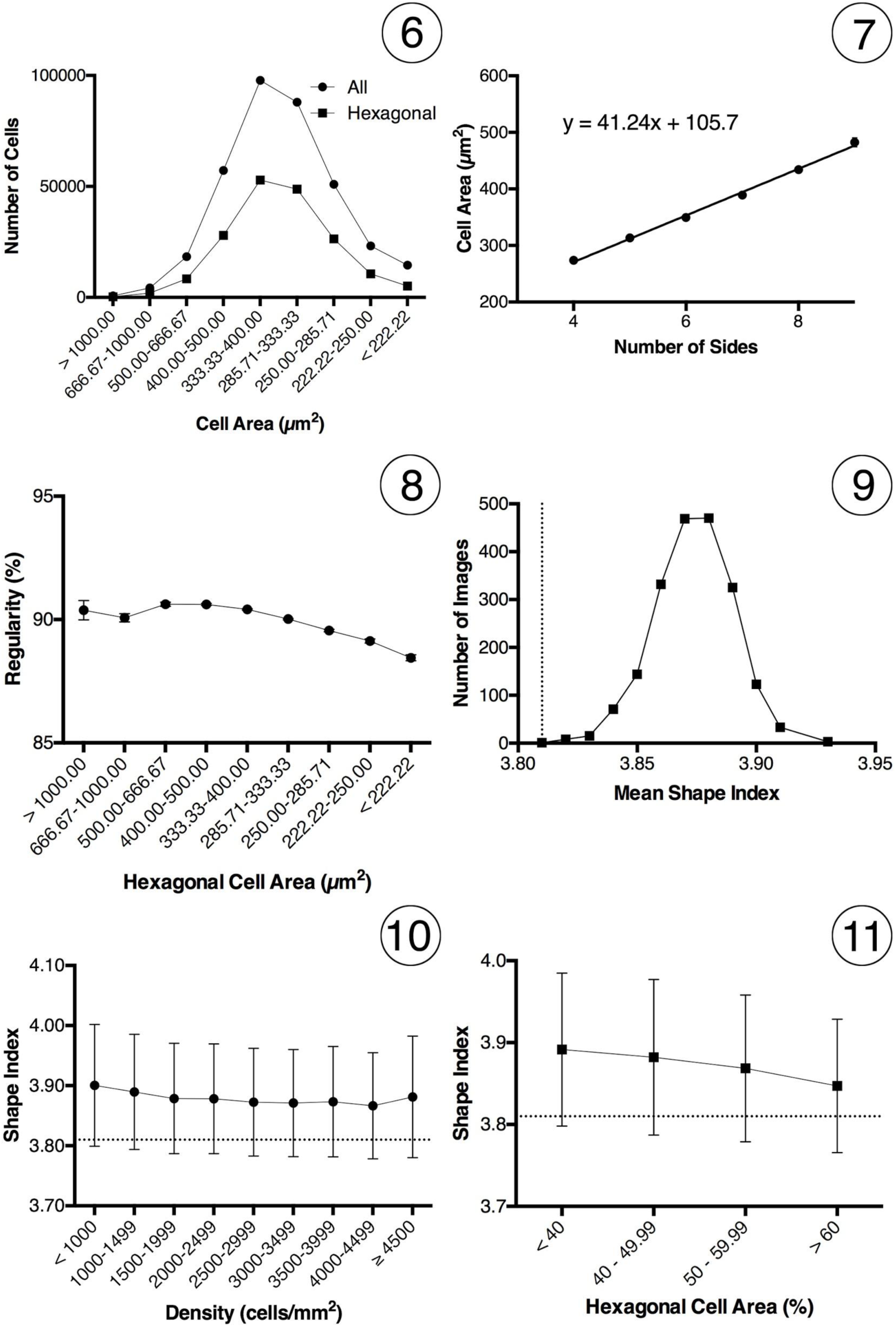
(6) Cell Area Distribution. (7) Cell Area *vs.* Number of Cell Sides (with linear regression line). (8) Hexagonal Cell Area *vs.*% Hexagonal Cell Regularity. (9) Image Shape Index Distribution. (10) Cell Shape Index *vs.* Image ECD. (11) Hexagonal Cell Shape Index *vs.*% Image Hexagonal Cell Area. Graphs 7, 8: Mean ± SEM. Graphs 10-11: Mean ± SD. Graphs 9-11: dotted line = CJB.

As expected, hexagonal cells dominated the endothelial mosaics across the full ECD range. The proportional area contribution of hexagonal cells slowly increased with density (Graph 2), and simultaneously the areas of pentagonal, heptagonal and octagonal cells slowly decreased. There was a constant background population of rectangular cells comprising between 1-1.3% of the area, which also decreased in area as the density increased. The largely constant relative proportions of different polygons through the bulk of the ECD range shows self-similar topological coarsening [Cantat *et al.*, 2013]. The increased proportional area of heptagons and octagons over rectangles and pentagons seen here may be partly due to the negative curvature of the posterior corneal surface [Weaire and Rivier, 1984], as well as the Lewis law.

Pleomorphic composition in proliferating tissues has been modelled to show that cell division processes alone will intrinsically converge on an irregular but predictable polygon distribution regardless of initial composition (28.9% pentagons, 46.4% hexagons, 20.8% heptagons, and lesser numbers of other polygon types) [Gibson *et al*., 2006; Nagpal *et al.*, 2008]. The corneal endothelium is the opposite of a proliferating tissue, where cells instead progressively die without replacement. While polygon proportions found in this study had some similarities to the model at low densities, there were quite different compositions throughout the bulk of the density range, suggesting that other factors must also contribute.

This notable stability in the number of hexagonal cells over the bulk of the range of ECD is predicted by both the Euler formula and the von Neumann-Mullins relation (Graph 4) [Cantat *et al*., 2013; Sánchez-Gutiérrez *et al.*, 2016]. As found elsewhere [Lewis, 1926; Nagpal *et al*., 2008], even though the average number of cell sides was six there was still considerable diversity, with approximately 5 different cell shapes per image over the bulk of the ECD range (Graph 3). The average values plotted in Graph 4 additionally show that the mean topological charge was zero over the bulk of the ECD range in the corneal endothelium.

While the simplest measure of polymegathism is the mean and spread of the percentage areas of cells in the mosaic, this provides no information about the regularity of individual cells [Doughty, 1992]. Another measure is the CV of the cell area, but due to the inherent ambiguity in interpreting this its use has been discouraged [Doughty, 1989; Doughty, 1990]. As discussed previously, the *ex vivo* cell area CV values found in this study were lower than those reported in many other mostly *in vivo* studies, probably reflecting that the VD ‘cell’ shapes are composed of straight lines rather than the tortuous curves generated by edge-based cell segmentation techniques [Brookes, 2017a]. However, cell area CV values in the range reported here have been seen in studies of foetal and children’s corneas [Farhan *et al*., 2014; Ko *et al*., 2001; Müller *et al*., 2000].

This current study has applied the CV to cell shape as well as size in order to further understand the complex topology of the endothelial mosaic. Both the mean cell size and shape CV decreased as ECD increased (Graph 5). The CV of number of sides has not been studied previously and showed that smaller cells (higher ECD) have less variation in shape as well as size.

Doughty introduced analysis of the area-side relationship as a means of comparing different endothelia that removes ambiguities inherent in the CV of cell area measurement [Doughty, 1990] and found a linear relationship between the cell area and the number of sides for cells with 4 - 8 sides [38,60]. This linear relationship was confirmed here using a far larger dataset (Graph 7), which fits the data extremely well extending to at least 9-sided cells, conforming to the Lewis law [Chiu, 1995].

As discussed previously, this study defined hexagonal cell regularity in a novel way [Brookes, 2017a], and while it was notably stable around 90% regular over the bulk of the cell size range, there was a decreasing trend where small cells had the least regularity, from a plateau around 400-500μm^2^(Graph 8). The hexagon has a unique geometry where the length of each side is equal to the centre-vertex radius, balancing both internal and surface forces, and making it the most optimal configuration both geometrically and thermodynamically [Lecuit and Lenne, 2007; Honda, 1983; Rao *et al*., 1982; Yee *et al*., 1985].

### 4.3 Shape Index and Tissue Phase

The most recent SI analyses have all been performed using either computer modelling or *in vitro* cell culture with a range of animal cell lines [Bi *et al.*, 2016; Park *et al*., 2016], so this study may be the first analysis of SI and the CJB in *ex vivo* human tissue.

The distribution of mean SI values in the endothelial images peaked at a mean (and median) of 3.87 (Graph 9). All but one of the 2000 images had a mean SI greater than the CJB at a SI of 3.81, showing that the endothelia were predominantly fluid. This was confirmed in Graph 10 across the entire ECD range, though the mean SI was largely independent of ECD with a very small decreasing trend. The SD plotted in Graph 10 shows that the SI of the cells comprising these endothelia spanned both sides of the CJB across the entire ECD range. The SI was however dependent on the percentage area of hexagonal cells in the image, where the higher the percentage of hexagonal cells the closer they were located to the CJB (Graph 11). Again, these hexagonal cell regions were comprised of cells with a range of SI’s. The relative proportions of those below the CJB increased along with the proportional hexagonal cell area.

At the cell level (Graph 12), the SI showed a bimodal distribution that was perfectly centred at the CJB. The largest proportion of cells had SI’s greater than this boundary, confirming that endothelia dominated by them would be in a fluid phase. A smaller cell peak at SI’s less than the boundary indicates a smaller cell population comprising solid endothelia. The shoulder in the graph at the CJB suggests some form of phase switching via shape transformation across this energy barrier, mostly toward more fluid mosaics.

**Graphs 12-17.**
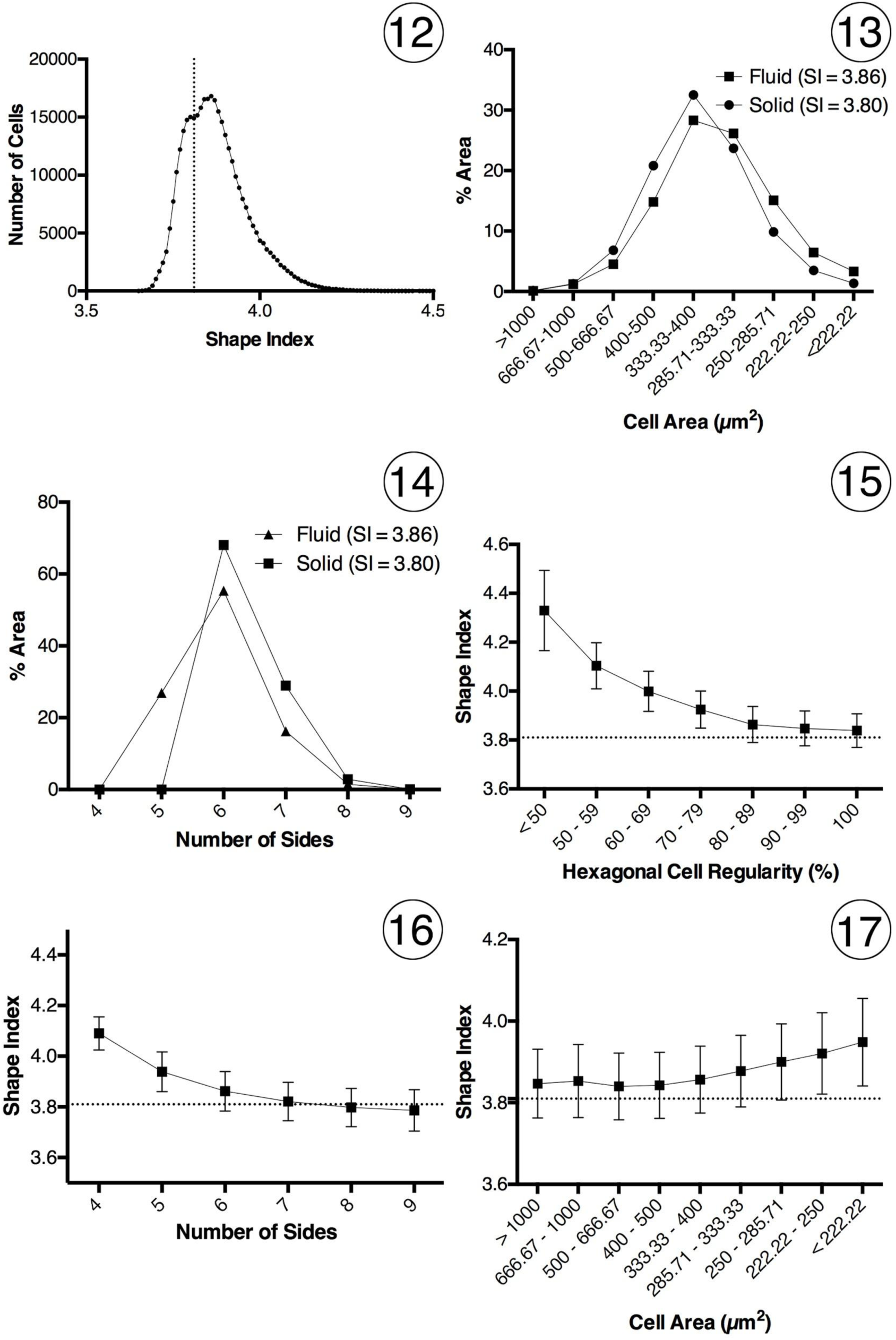
(12) Cell Shape Index Distribution. (13) Proportional areas of cells adjacent to the CJB. (14) Proportional cell areas of pleomorphs adjacent to the CJB. (15) Hexagonal Cell Regularity *vs.* Cell Shape Index. (16) Number of Cell Sides *vs.* Cell Shape Index. (17) Cell Area *vs.* Cell Shape Index. Graphs 15-17: Mean ± SD. Graphs 12, 15-17: dotted line = CJB.

Analysis of the cell populations under the two SI data points at the peaks of the bimodal distribution either side of the CJB at 3.80 (solid) and 3.86 (fluid) revealed quite different characteristics, as summarized in Table 1 and Graphs 13-14. Cells in the solid region contained a greater proportion of larger cells with six or more sides. The hexagonal cells were also more regularly shaped. There were no cells in this region with less than six sides. The dynamic balance of these mixtures suggests that shape and size change of only a small proportion of cells could radically modify the phase properties of the cell collective, and that this diverse morphological composition may be integral for the maintenance of endothelial structure and function.

SI was also dependent on hexagonal cell regularity (Graph 15), where the most regular hexagonal cells were closest to the CJB. The mean SI skated asymptotically toward the CJB, confirming that most endothelia were in a fluid phase, where rearranging their mosaics requires little energy. Beyond a regularity of about 80% the spread of values shows a proportion of cells crossing over the CJB. Hexagonal cell properties are uniquely balanced in relation to the CJB, where small individual morphometric changes will have large effects on collective cell phase. The bottom of the range found in the dataset corresponded to the SI of a perfectly regular hexagon (3.72), well into the solid side of the boundary.

The SI was also dependent on both the number of cell sides (Graph 16) and cell area (Graph 17). Only cells with less than 6 sides had SI’s above the CJB. While the mean hexagonal cell SI was also above the CJB, a proportion of these cells had SI’s below it. Mean heptagonal cell SI was located very close to the CJB (3.82), and cells with more than seven sides had mean SI’s below the CJB. However, in all but the two ten-sided cells in the dataset there was still a proportion with SI’s on the fluid side of the CJB. While the smallest cells had the largest SI’s, this trend was lost than when averaged out at the image level, where individual endothelia were comprised of cells with a range of SI’s (Graph 10).

### 4.4 Centroidal Voronoi Tessellation

This study computed the VD from a point cloud of cell centroids so was a form of Centroidal Voronoi Tessellation (CVT) [Du *et al*., 1999]. An optimized CVT topologically relaxes toward a uniform spatial distribution of centroids [Du *et al*., 2006] and applying Gersho’s conjecture to a CVT with a large number of centroids shows it tends asymptotically toward comprising of only regular hexagons [Du and Wang, 2005]. This aligns with the Euler formula, where endothelia will tend to be dominated by regular hexagonal cells. Graphs 11 and 15-16 show that this arrangement shifts the cell collective closer to a solid phase.

The proportional polymorph area can also be compared to an idealized curve of all probable size-shape relationships using a CVT pathway graph [Sánchez-Gutiérrez *et al*., 2016]. This hypothetical framework can be used to compare the organization of a wide range of two-dimensional tissue architectures and suggests that they cannot present infinite organizations but are constrained to certain combinations of polygon distributions. Comparison of the data from this study with plots published by Sánchez-Gutiérrez *et al*. indicates that the *ex vivo* corneal mosaics are not fully optimized CVT’s (data not shown) [Sánchez-Gutiérrez *et al*., 2016]. Such deviations from these idealized plots imply physical cellular constraints modifying the balance of forces within the tissue due to genetic or disease factors, thereby limiting tissue organization.

### 4.5 Conclusion

The data presented here shows that collectively the corneal endothelial mosaic acts as a glassy fluid, governed by well-characterized physical laws. The tissue is a form of viscous foam, with a cytoskeleton and intercellular junctional complexes tightly anchored to Descemet’s membrane. Individual cell death transiently and locally increases collective cell entropy, so the endothelium must expend energy in order to rearrange and maintain function. Independent of ECD, this organization progressively tends toward a CVT with minimum free energy, comprised of a greater proportion of more uniformly sized cells with six or more sides. Becoming simultaneously more ordered and solid, each cell requires more energy to move. This rigidity is an emergent property that efficiently locks cells in place and maintains endothelial function. *In vivo*, this homeostasis is opposed by other factors such as age, environment, genetics, and pathology.

